# Microbial Dysbiosis and polyamine metabolism as predictive markers for early detection of pancreatic cancer

**DOI:** 10.1101/342634

**Authors:** Roberto Mendez, Kousik Kesh, Nivedita Arora, Leá Di Martino, Florencia McAllister, Nipun Merchant, Sulagna Banerjee, Santanu Banerjee

**Author notes:** Contributed equally (Roberto Mendez, Kousik Kesh). Funding information:* This study was partially funded by NHLBI R21 HL125021 to Santanu Banerjee, NCI R01CA184274 to Sulagna Banerjee and University of Miami Institutional support to both. The funders had no role in study design, collection, analysis or interpretation of data.

## Abstract

**Purpose:** The lack of tools for early detection of pancreatic ductal adenocarcinoma (PDAC) is directly correlated to the abysmal survival rate in patients. In addition to several potential detection tools under active investigation, we tested the gut microbiome and its metabolic complement as one of the earliest detection tools that could be useful in patients at high-risk for PDAC.

**Experimental Design:** A combination of 16s pyrosequencing and whole-genome sequencing of gut microbiota was used in a spontaneous genetically engineered PDAC murine model (KRAS^G12D^TP53^R172H^Pdx^Cre^ or KPC). Metabolic reconstruction of microbiome was done using the HUmanN2 pipeline. Serum polyamine levels were measured from murine and patient samples using standard methods.

**Results:** Results showed a progressive Proteobacterial and Firmicutes dominance in gut microbiota in early stages of PDAC development. Upon *in silico* reconstruction of active metabolic pathways within the altered microbial flora, polyamine and nucleotide biosynthetic pathways were significantly elevated. These metabolic products are known to be actively assimilated by the host and eventually utilized by rapidly dividing cells for proliferation validating their importance in the context of tumorigenesis. In KPC mice, as well as PDAC patients, we show significantly elevated serum polyamine concentration. Therefore, at the early stages of tumorigenesis, the gut microbial composition changes in a way to release metabolites that foster host tumorigenesis, thereby fulfilling the ‘vicious cycle hypothesis’ of the role of the microbiome in health and disease states.

**Conclusions:** Our results provide a potential, precise, non-invasive tool for early detection of PDAC, which will result in improved outcomes.

**Synopsis:** Gut microbiota changes during early stages of pancreatic ductal adenocarcinoma (PDAC) progression and contributes towards host polyamine pool. Both changes can be used as an early predictive marker for PDAC.

**Translational Relevance:** Pancreatic carcinogenesis progresses through pre-cancerous PanIN lesions to invasive cancer. Even though these morphological changes are histologically distinct, imaging techniques are not able to distinguish the pre-invasive PanINs from normal pancreas, making detection of a tumor at a precancerous stage impossible. Thus, majority of cases (85–90%) present with advanced pancreatic cancer at the time of diagnosis. This contributes to the dismal survival rate in this disease. Our study of gut microbiome analysis on KPC mice during tumor progression followed by metabolic reconstruction and experimental validation in human samples indicate that gut-microbiome analysis along with an analysis of the microbial metabolites can be developed as potential biomarkers for detection of PDAC at early stages when histological changes are not yet grossly apparent.

## Introduction

Pancreatic cancer is the 3^rd^ most common cause of cancer related deaths in United States with a very poor 5-year survival rate of 9%. The disease is characterized by relatively late onset of symptoms and rapid progression with very limited treatment options. Lack of efficient biomarkers that can facilitate early detection of the disease is one of the primary challenges of the field. With effective tools of early detection, the survival rate would eventually increase, as more patients would be able to have their tumors resected (1). Thus, there is an urgent need for non-invasive, discriminatory biomarkers for early detection of this disease. During carcinogenesis, a pancreatic tumor progresses through the pre-cancerous pancreatic intraepithelial neoplasia (PanINs) lesions (2,3). However, even though these changes in the pancreas morphology are histologically distinct, imaging techniques used in the clinics are not able to distinguish the early PanINs from the normal pancreas. As a result, detection of a tumor at an early stage becomes difficult. Retrospective study has shown that 20-25% of patients with pancreatic cancer develop diabetes mellitus 6-36 months before pancreatic cancer is diagnosed (4). This observation is also being evaluated as a possible indication of early detection in this disease.

Pancreatic tumorigenesis is heavily dependent on inflammation. Recent studies have shown that systemic inflammation plays a significant role in altering the microbial flora in the gut, leading to microbial dysbiosis (5,6). The role of gut microbiome, and specifically its dysbiosis in cancer development, is becoming more evident in recent years. It becomes increasingly apparent that the gut microbiome plays an integral role in modulating host physiology, including disease-specific unique cellular metabolism and immune function (7). Initial studies of the gut microbiome focused primarily on colon cancer, however, recent studies have shown that microbial changes are associated with development of melanoma (8), gastric cancer (9), lung cancer (10) as well as PDAC (11–18). It is now well-established that the gut bacteria play an integral role in cellular metabolism and immune function that become deregulated during carcinogenesis. While microbiome research has concentrated on its role in tumor progression, change in the microbiome and their metabolites have not been exploited as potential biomarker for early detection in pancreatic cancer (19–22).

Previous reports have suggested that oral microbiome dysbiosis occurs during pancreatic tumorigenesis (12,23). Bacterial species have been detected in pancreatic cysts, as well as in PDAC tumors (24). In addition, other studies have revealed that the microbiome plays an active role in stromal modulation (16) and depletion of the microbiome plays a profound role in reduction of pancreatic tumor burden (18).

Most of these studies, however, have utilized the most common sequencing approach of an amplicon analysis of the 16S ribosomal RNA (rRNA) gene (25,26). This method is limited by the fact that the annotation is based on putative association of the 16S rRNA gene with an operational taxonomic unit (OTU). Since OTUs are analyzed at the phyla or genera level, the study is often unable to detect species within those taxa. An alternative approach to the 16S rRNA amplicon sequencing method is whole genome shotgun sequencing (WGS) which uses sequencing with random primers to sequence overlapping regions of a genome allowing for the taxa to be more accurately defined at the species level (27).

In the current study, we analyzed the microbiome of in a genetic mouse model for PDAC (KRAS^G12D^TP53^R172H^Pdx^Cre^ or KPC) and age-matched controls using WGS at very early time points of tumorigenesis. During these time points, the KPC mice do not show any detectable tumors in their pancreas. Our results show that at these early time points, the histological changes in the pancreas correspond to a significant change in certain gut microbial population. Our predictive metabolomic analysis on the identified bacterial species reveal that the primary microbial metabolites involved in progression and development of PDAC tumors are involved in polyamine metabolism. Consistent with our analysis in mice, estimation of polyamine from serum of PDAC patients show that total polyamine concentration in increased in PDAC patients compared to healthy volunteers. Furthermore, serum polyamine levels in KPC mice show an increased concentration of polyamines as tumors progress from PanINs to PDAC. Together these observations indicate that analysis of gut microbial flora along with an analysis of the microbial metabolites can be developed as potential biomarkers for detection of PDAC at early stages when histological changes are not yet grossly apparent.

## Results

### Gut microbial profile in KPC mice diverges from age-matched control mice over time

We employed a variety of indices to understand the composition and distribution of bacterial communities (species richness, evenness, distribution; alpha-diversity) in one and six months old KPC; and age-matched control mice. While controversies exist in the implementation of various diversity indices on 16s microbial data, originally developed for ecological studies with macro-organisms, a combination of these indices for 16s phylogenetic studies provide an overall spectrum of the composition of each group (28). We started with the Shannon’s H index (29), which accounts for both the abundance and evenness (measure of species diversity and distribution in a community) in both age groups and genotypes (**Figure 1A**). The Shannon’s H index showed no difference between the overall composition of the two age groups (1 and 6 months old control or KPC mice), implying that the numbers of different OTUs encountered (diversity), as well as the instances these unique OTUs were sampled (evenness) was relatively similar between the two groups. Since Shannon’s index is blinded to the identities of the OTUs, we also included Chao1 diversity (30) which accounts for the rarity or abundance of individual OTUs and provides a measure of species richness for individual groups which can then be compared (**Figure 1B**). The Chao1 diversity analysis showed significantly diminished OTU richness in KPC mice compared to control mice at 6 months of age. Faith’s phylogenetic diversity (PD_whole_tree) represents taxon richness of each sample expressed as the numbers of phylogenetic tree branches encountered for each sample based on sequenced OTUs (31). In a similar manner as was seen in the Chao1 index, a significant reduction in OTU diversity was observed in KPC mice compared to control in 6-month age groups (**Figure 1C**).

**Figure 1:**
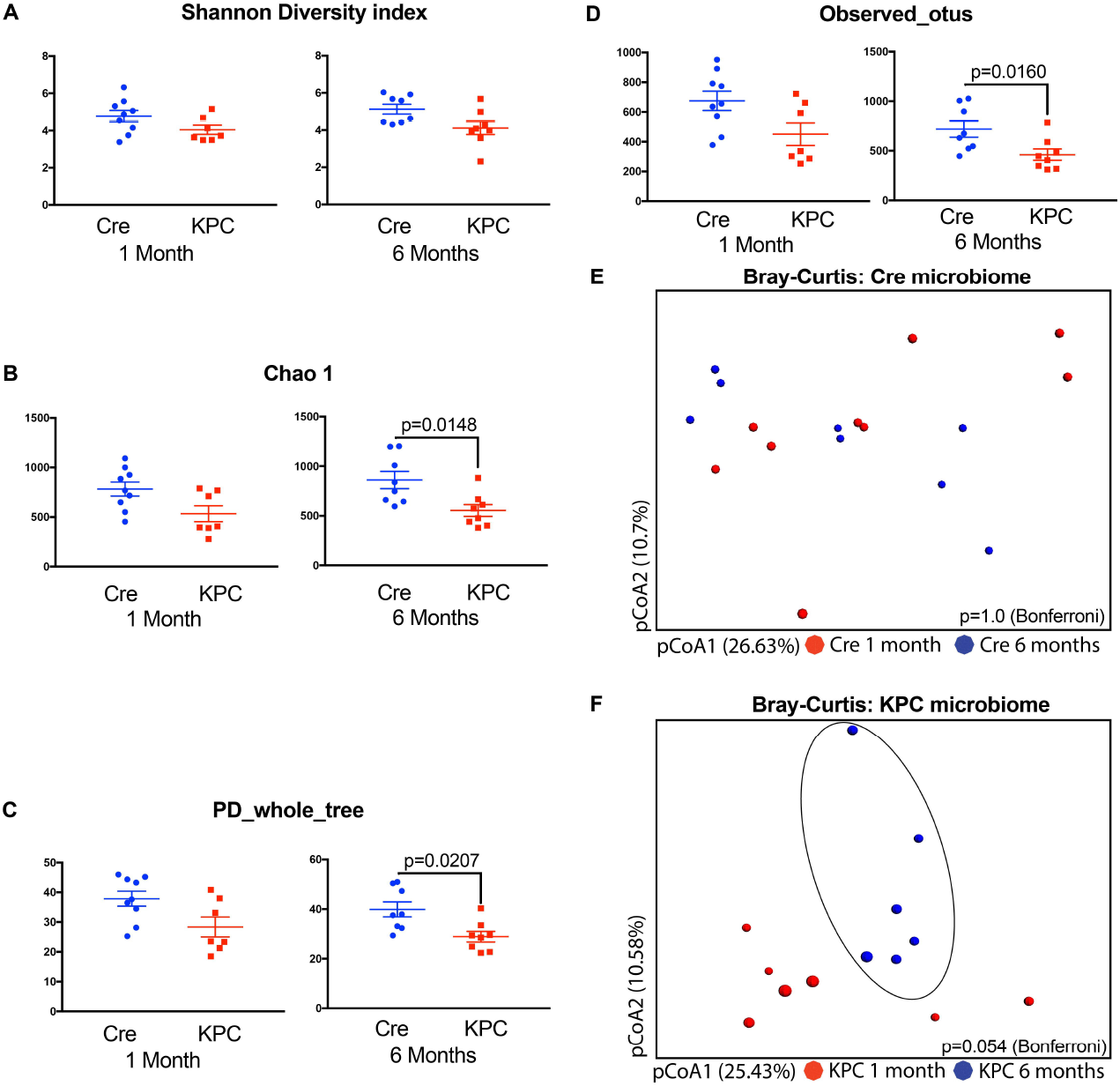
KPC microbiome exhibits clustering between 1 and 6 months, compared to control microbiome with 16s pyrosequencing. Various alpha-diversity indices were measured for the two genotypes and two time-points. **(A)** Shannon’s H index was similar between the age groups and genotypes, whereas **(B)** Chao1 index, **(C)** Faith’s Phylogenetic Diversity and **(D)** Observed OTU indices were significantly down in 6 months old KPC animals, compared to age-matched control animals. Bray-Curtis Principal Co-ordinate Analysis showed that **(E)** control animals did not form distinct clusters between 1 and 6 months of age, whereas **(F)** KPC animals for a distinct cluster at 6 months age, compared to 1 month old animals. This difference, however, was not statistically significant (p= 0.056) with 2-tailed test of significance with Bonferroni correction [n= 6 to 8 per group].

The disparity between Shannon index and Chao1/PD can be described through the ‘Observed OTU’ metric (**Figure 1D**), which accounts for an absolute presence/absence of the OTU irrespective of the number of instances it is represented in a sample. This index also exhibited a significant reduction of richness in 6-month-old KPC mice compared to the control animals. This disparity implies that some species of microbes are actually getting diminished with tumor development. The evenness of distribution makes the Shannon index comparable between the two age groups, despite differences in alpha-diversity using other matrices. This is not uncommon, as described in other microbial studies(32), and ensures that the differences in the beta-diversity (difference between groups) are not entirely due to differences in absolute OTU counts (32).

Next, we used Bray-Curtis Principle Co-ordinate Analysis (pCoA; beta-diversity) to qualitatively examine differences in microbial composition and associated changes due to age in control (**Figure 1E**) and KPC mice (**Figure 1F**). Microbial composition of control mice did not change over the course of 6 months, whereas a tendency towards clustering was observed in KPC 6 months old mice. However, the differences were statistically insignificant (p=0.054). Since there was a significant drop in phylogeny-informed diversity indices for KPC mice over time, we decided to study the major differences between the two KPC age-groups, in the context of identified genera, family and class of microbes.

### There are significant early differences in a small group of microbes at the class and genera level between one- and six-month old KPC animals

An analysis of the five major phyla revealed a relatively unchanged Bacteroides and a trend towards decrease in relative abundance in Actinobacteria, Deferribacteres, Firmicutes and Proteobacteria (**Figure 2A**). None of these phyla had statistically significant changes over time. However, when we looked at the class-level in the hierarchy, several classes of bacteria showed significant changes in relative abundance between 1-month- and 6-month-old KPC animals (**Figure 2B**). Compared to non-significant changes in class Bacteroidea, there was a significant drop in the relative abundance for classes Clostridia, Bacilli, Erysipelotrichia (All phylum Firmicutes), Deferribacteres (Phylum Deferribacteres), class Actinobacteria (Phylum Actinobacteria) and Delta-, Epsilon-, Gamma-, and Betaproteobacteria (All phylum Proteobacteria). Within Phylum Proteobacteria, however, there was a significant relative expansion of class Alphaproteobacteria in 6-month-old KPC animals. At the genus level, 6 genera exhibited significant relative expansion, whereas 19 showed diminished relative abundance from 1 to 6 months in KPC animals (**Figure 2C**). Interestingly, it is obvious from **Figure 2c** that there is a clear disconnect between the trends seen in upper hierarchical levels and genera-level changes, specifically for genera showing relative expansion (1–6, **Figure 2C**). While Actinobacteria as a phylum (**Figure 2A**) and class (**Figure 2B**) are seen to have diminished relative abundance, *Adlercreutzia* has a significant relative expansion in 6-month-old KPC animals. Similarly, within phylum Firmicutes, classes Erysipelotrichi, Clostridia and Bacilli exhibit lower relative abundance (**Figure 2B**), genera within the classes respectively, *Turicibacter, Coprococcus* and *Staphylococcus* are significantly elevated in 6 months old KPC animals. In a similar vein, genera including *Bacteroides, Parabacteroides* and *Prevotella* (all Phylum Bacteroides) exhibit significantly lower abundance in 6-month-old KPC animals, despite no differences in the Phylum or Class levels of their hierarchy. Furthermore, upon mapping all independent OTUs (n=24) mapping to genus *Bacteroides*, several minor OTUs also have significantly elevated abundance in 6 months old KPC animals (**Supplementary Figure 1**) but are beyond the scope for any further identification due to the limitations of the 16s platform used. For mapping, microbial changes in spontaneous PDAC model and predicting functional attributes, it became abundantly clear that a better understanding of the early changes in genus/species level is of paramount importance. Unfortunately, phylogeny based on 16s amplicon sequencing does not provide sufficient genus/species level coverage, and, hence, we decided to restrict the time points and employ shotgun metagenomics for further studies.

**Figure 2:**
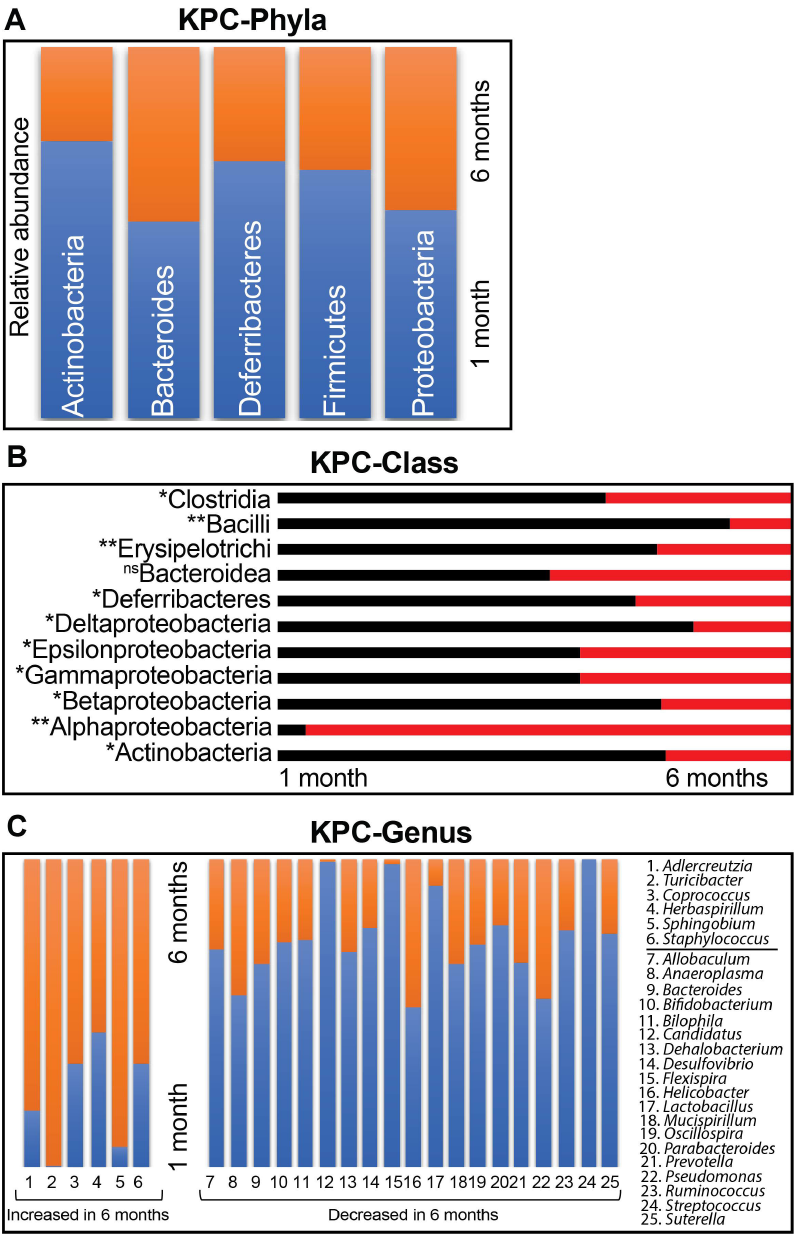
KPC animals show significant differences at the Class and Genera levels. In a head-to-head comparison between the OTUs representing the five major phyla, **(A)** Bacteroidetes did not show any difference between 1 and 6 months old KPC mice. The other four phyla showed reduced relative abundance, which were statistically insignificant. At the Class level **(B)** however, Alphaproteobacteria exhibited significantly high relative abundance in 6 months old KPC. With Bacteroidea unchanged between the two age groups, all other Classes exhibited diminished relative abundance in 6 months old KPC mice [Test of significance-non-parametric Mann-Whitney U test. *p<0.05, **p<0.01]. At the Genus level **(C)**, six genera showed significant increase in relative abundance from 1 to 6 months of age, compensated by 19 genera with severely diminished relative abundance at the same time [All genera-p<0.05].

### Microbial Dysbiosis occurred during early processes of tumor development

For the pre-neoplastic time points, in order to obtain functionally relevant data with species-level identification, we performed whole genome sequencing on control and KPC animals at 2, 3, and 4 months of age in animals that did not have visible tumors. A total of 1359 genera were identified with 100% species-level coverage. Initial comparison of the overall microbial composition between control and KPC mice showed progressive microbial changes in the KPC mice from 2 to 4 months (**Figure 3A-3C**). At 2 months of age, microbiome of control animals and majority of KPC animals form a single cluster (circled), with apparent changes in some of the KPC animals (**Figure 3A**). By the third month of age, control animals maintained the cluster, one KPC animal shifted out of the circled cluster, accompanied by mortality of another KPC mice, which was outside the cluster from 2 months of age (**Figure 3B**). Finally, at 4 months age, all KPC animals except one, were seen to acquire a distinct microbial composition compared to the control animals (**Figure 3C**). At this point however, all KPC animals outside the cluster at 3 months of age had not survived. This phenomenon is also evident from the Class/genus level heatmap of top 50% representative bacteria at 2- and 4-month-old control and KPC mice, where we do not see any distinct pattern between the two genotypes at 2 months age (**Supplementary Figure 2**), while a distinct pattern can be seen emerging in 3 out of 4 KPC animals at 4 months of age, consistent with the pCoA findings in **Figure 3C** (**Supplementary Figure 3**). In general, the KPC mice seem to maintain a higher species richness and evenness of distribution, which is significantly higher in the 3^rd^ and 4^th^ months of age (**Figure 3D**).

**Figure 3:**
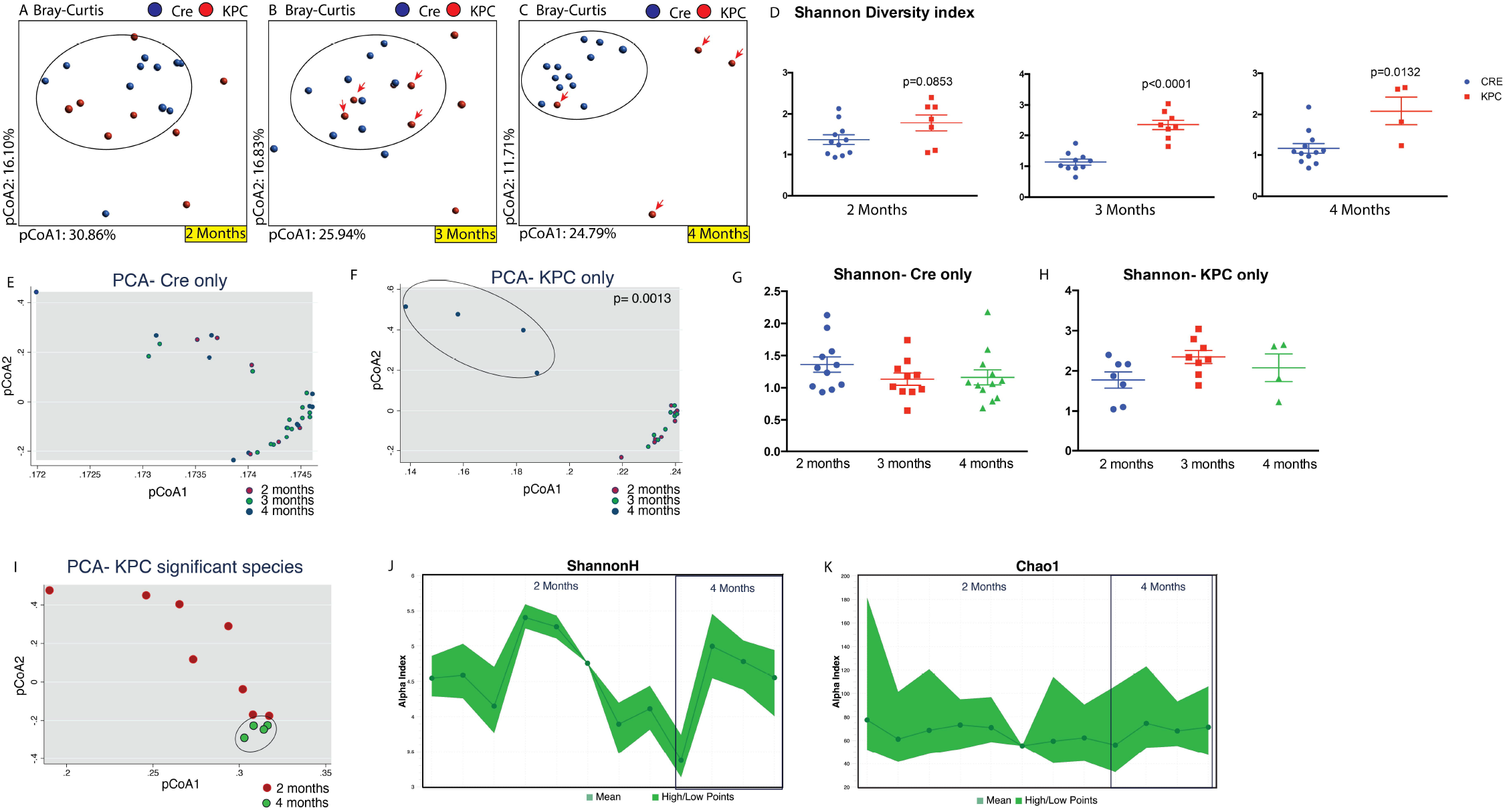
Whole genome sequencing of control and KPC mice at ages 2, 3, and 4 months. As seen above, the microbial composition of KPC animals, which is similar to control (circled) animals at 2 months age **(A)**, starts changing by 3 months age **(B)**. By 4 months, there are significant differences in the control and KPC microbiome **(C)**. In this experiment, only 4 KPC animals survived for 4 months (arrows point to the individual animals which survived from 3^rd^ to 4^th^ month). This is accompanied by significantly increased Shannon’s H alpha-diversity index for 3- and 4-month old KPC animals, compared to their control counterparts **(D)**. While control animals do not exhibit change in microbial composition in pCoA plot with age **(E)**, four surviving members of 4-month-old KPC animals were seen to cluster separately from the 2 or 3 months of age collections **(F)**. The Shannon index was not different within control **(G)** or KPC **(H)** animals over time. In KPC animals, between 2 and 4 months age, 82 species were found to be significantly different (see Supplementary Figure 4). Analysis of those 82 species with pCoA plot shows tight clustering of 4-month-old animals, compared to when they were 2 months old **(I)**. Neither the Shannon **(J)** or Chao 1 **(K)** indices were different for the two age groups, when analyzed for the significantly changing species only.

This prompted us to analyze the control and KPC microbiome separately across all three months.

The microbiota of control animals, as evident from **Figure 3A-3C**, does not alter from the 2^nd^ to the 4^th^ month (**Figure 3E**). While there is some variability among individual animals across the time points, there were no distinct clusters within the defined time points. In KPC animals however, the microbiome at 4 months age was distinctly and significantly different from the other age groups (**Figure 3F**). As mentioned earlier in **Figure 3D**, KPC microbiome maintains a higher level of species richness and evenness. However, there were no significant changes in control (**Figure 3G**) or KPC (**Figure 3H**) mice across the time points measured. Hence, the differences in the microbial composition with age is not influenced by the richness/evenness component of evaluation.

Next, we concentrated on the significantly changed species within the 2 months and 4 months KPC animals (**Supplementary Figure 4A**). A total of 89 species, across all classes and phyla, were found to be significantly up-or down-regulated with age. Among the significantly changing species, only 7 were found to lose abundance, with 4 belonging to phylum Firmicutes and 3 from Proteobacteria (all gammaproteobacteria). The remaining 82 species exhibited significantly expanded relative abundance from 2 months to 4 months of age, represented by 14 different bacterial phyla (**Supplementary Figure 4B**). As shown, approximately 40% of these species belong to Phylum Proteobacteria, followed by Firmicutes (18%), Actinobacteria, Bacteroidetes, Cyanobacteria and Euryarchaeota (all 6-7%). The significantly altered species in 4 months old KPC animals were compositionally close and formed a tight cluster in a pCoA plot, compared to 2 months old KPC animals (**Figure 3I**). This is despite no significant differences seen in the Shannon (**Figure 3J**) or Chao1 (**Figure 3K**) indices, implying that the relative abundances of the significantly changing species in 4-month-old KPC animals are influenced by the disease and not the richness or evenness of the microbiome.

### The metabolic landscape of the microbiota shows a shift from dominance of energy metabolism to upregulation in polyamine and lipid metabolism with PDAC tumor progression

We then characterized the functional profile of the microbiota of 2-month-old and 4-month-old KPC mice using the HUMAnN2 pipeline (33,34). The resulting pathways were classified and curated manually into: (a) significant pathways in 2-month-old KPC only, (b) significant pathways in 4-month-old KPC only and (c) significantly changed pathways in 4-month-old KPC, compared to 2-month-old KPC animals (continuously valued relative abundance). We found that in 2 months old KPC, there is a distinct dominance of energy metabolism pathways in the 227 pathways identified (top 25 pathways depicted in **Figure 4A**). Within the top 25 pathways, apart from energy metabolism, pathways contributing to cell division (e.g. amino acid biosynthesis and nucleotide biosynthesis pathways) were adequately represented. However, in 4-month-old KPC animals, the landscape was found to be significantly dominated by polyamine biosynthesis (**Figure 4B**). Biosynthesis of polyamines Putrescine, Spermidine, and Spermine dominated the metabolic landscape of the dysbiotic microbiome of 4-month-old KPC mice. The rest of the pathways were represented by metabolic shunt pathways and lipid metabolic pathways. Next, we looked into the differential upregulation of pathways between 2^nd^ and 4^th^ month KPC animals (**Figure 4C**). We found a 29% upregulation of polyamine biosynthesis pathways in 4 months KPC animals, followed by 24% upregulation in nucleotide biosynthesis pathways. Essentially, more than 50% of bacterial metabolism seems to be promoting DNA synthesis/ replication pathways, where the metabolic products are readily assimilated by the host (35–37). This is supported by a 14% upregulation of lipid biosynthetic pathways, important for *de novo* cell membrane synthesis/remodeling.

**Figure 4:**
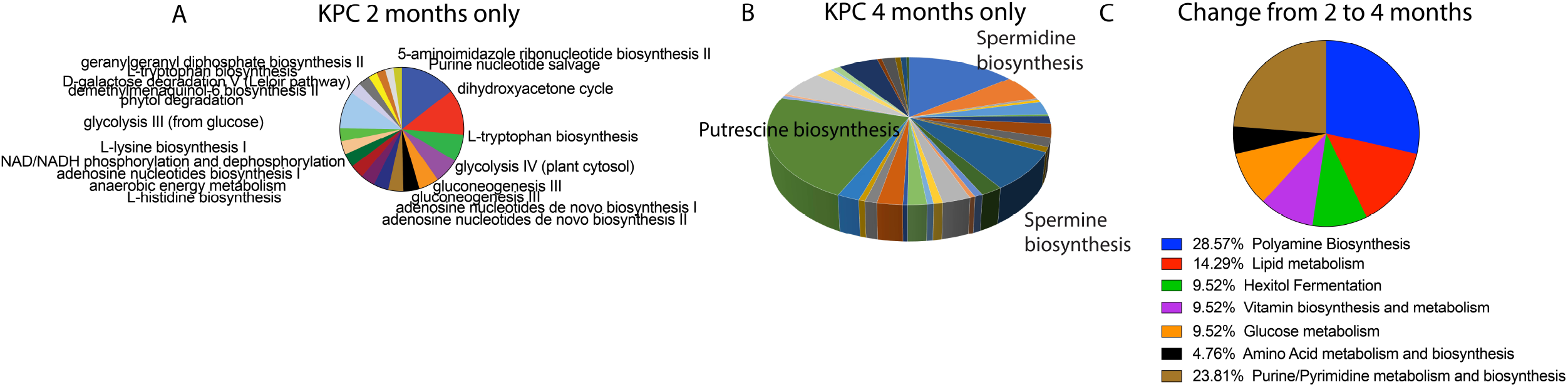
Metabolic reconstruction of the microbiome. The HUMAnN2 pipeline generated the pathway abundance list from whole genome sequencing input, and we manually curated it for pathways found in **(A)** 2-month-old KPC only and **(B)** 4-month-old KPC only. While 2 months only was dominated by energy metabolism pathways among others (see Table 1), 4 months only was dominated by Polyamine biosynthesis pathway. Overall, between the ages of 2 months and 4 months in KPC animals **(C)**, the most significant metabolic pathways were dominated by biosynthetic pathways, where the majority of metabolites are exchanged between the host and the microbiota.

To validate our predictive metabolomics, we next estimated polyamines from serum samples of KPC animals at different stages of tumor progression. Our results showed that while there was low polyamine in the serum of the animals of 1-2 month age, there was a significant increase in the serum concentration of these polyamines in 4 months animals that had PanIN2 and PanIN3 lesions but no observable tumor. Serum Polyamine concentration increased further in a full tumor (6 month and older) mice (**Figure 5A**). To further validate this, we next estimated polyamines in the serum from PDAC patients and compared their concentration to that in serum from normal healthy volunteers. Our results showed that in PDAC patients, serum polyamine concentrations were significantly increased (**Figure 5B**).

**Figure 5:**
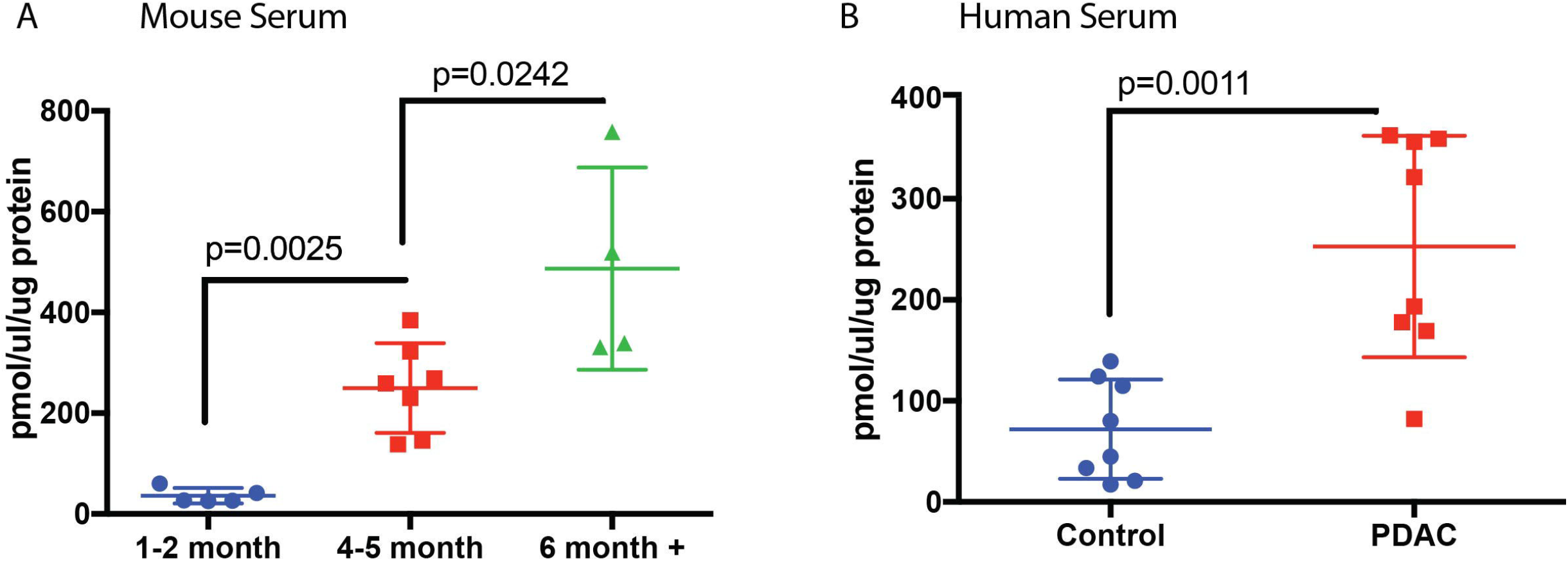
Serum polyamine levels are significantly high in spontaneous murine model of PDAC and in PDAC patients. When we measured the actual total polyamine levels in KPC mice serum (D), significant elevation was seen with progressing age and cancer. Similarly, serum polyamines were found to be significantly elevated in PDAC patients, compared to healthy controls (E). [n= 5-8 for mice; n= 8 for human serum samples]. Test of significance was 2-tailed, non-parametric Mann-Whitney U test. P-values are exact and mentioned in the figure.

**Figure 6:**
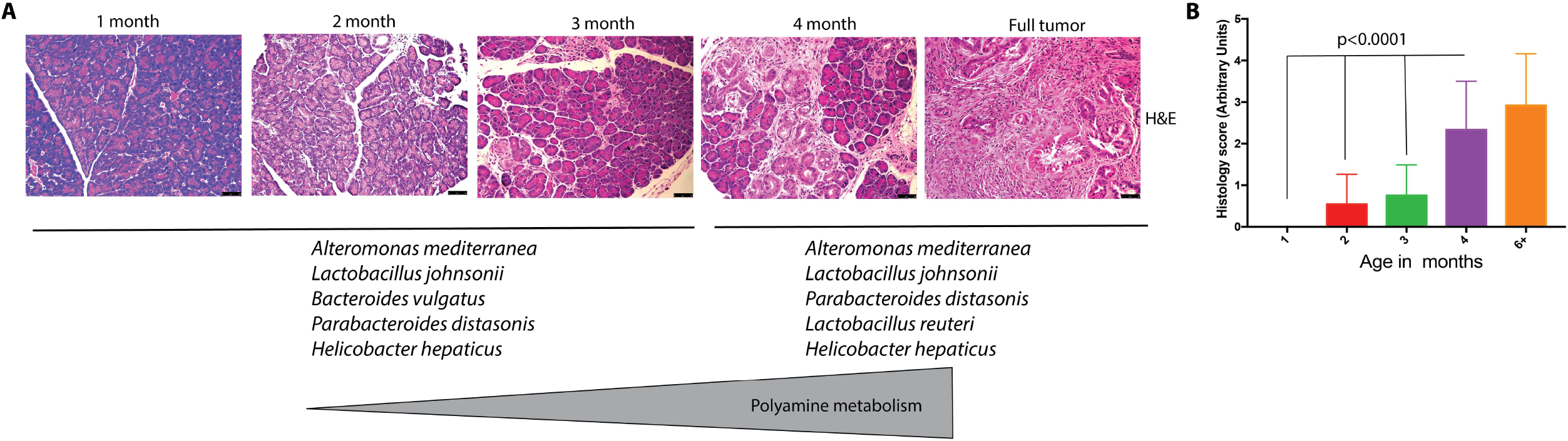
Progression to tumor and microbial/metabolic changes. With progressive cellular disorganization and tumor development in KPC mice (**A**; representative H&E staining), top bacterial species between 2 months and 4 months are joined by *Lactobacillus reuteri.* Lactobacilli is known to actively participate in polyamine metabolism. **(B)** Blinded histological scoring of PDAC progression in KPC mice [n=6/group with 4 fields per section]. Test of significance for (B) was non-parametric Mann-Whitney U test.

## Discussion

The role of microbiome in disease development, progression, and therapy are becoming increasingly clear. While this became apparent in colon cancer and other colonic disorders like inflammatory bowel disease where the bacterial population in the gut has direct influence on the inflammatory milieu of the disease, its influence in other diseases is only starting to be acknowledged (8,34,38). Changes in gut microbiome have now been associated with therapeutic response in melanoma (19,39), hepatocellular carcinoma, (14) and even in PDAC (16,18). While the current focus in microbiome research in PDAC has been primarily focused on therapy and its response with respect to microbial changes, there have been almost no studies on the role of this dysbiosis with respect to disease development and progression. Earlier studies have revealed that changes in the salivary and oral microbiome may correlate with risk of PDAC (12). In a recent study by Ren et al, a miSeq analysis showed that some potential pathogens like *Veillonella*, *Klebsiella*, and *Selenomonas* and LPS-producing bacteria including *Prevotella, Hallella*, and *Enterobacter* were enriched in pancreatic adenocarcinoma, whereas probiotics including *Bifidobacterium* and some butyrate-producing bacteria, such as *Coprococcus, Clostridium IV, Blautia, Flavonifractor* and *Anaerostipes* were reduced (40).

KPC mice have been a gold standard for understanding the progression of pancreatic tumors (41). In the current study, we found a distinct microbial dysbiosis between the microbiome of mice with no tumor at 1 month of age compared with mice with PDAC tumors at 6-months of age. To further resolve when the microbial dysbiosis occurs, we next collected and analyzed the fecal microbiome of 2-to 4-month old mice that did not have overt tumors (but had histological indication of panINs and some dysplasia). We gave an arbitrary dysplasia score in which histologically normal pancreases of one-month-old mice were considered 0 and histologic PDAC tumor-bearing mice were considered 4. Our results showed that microbial dysbiosis predominantly occurred at 4 months of age (mean histological score of 2.33), when there were no observable tumors in the animals. Our study corroborated the recent study by Pushalker et al.(16) We observed that there is a distinct difference in the microbial population as a pancreatic tumor develops (**Figure 1**). At this stage, a significant difference was observed at the class level, particularly for classes like Clostridia, Bacilli and Erysipelotrichi, Actinobacteria, and Deferribacteres (**Figure 2**). Interestingly, when we further analyzed the microbiome of KPC mice at an early “pre-tumor” stage in which there was no observable tumor (only histological indication), the microbial population seemed to emerge as a distinct pattern (**Figure 3**). The KPC animals appeared to maintain a higher species richness and evenness of distribution. Our analysis further showed that the significantly altered species in 4-month-old KPC animals were compositionally close and formed a tight cluster in a principal coordinate analysis despite no significant difference in the Shannon or Chao1 index, indicating that this difference was due to the progression of the disease (**Figure 3**).

Microbial population is known to influence the host metabolism in a number of ways. Studies have shown that bile acid metabolism is one of the main metabolic pathways that are affected by microbial dysbiosis (42). Similarly, short chain fatty acids that the exclusive products of microbial metabolism are known to affect the host epigenetic machinery (43). Our metagenomics analysis on the predictive metabolome of the microbial species shows that there is a distinct dominance of energy metabolism pathways (**Figure 4**). Interestingly, the major metabolic pathways that are deregulated from 2 months to 4 months of age in KPC mice are those involved in polyamine biosynthesis and pyrimidine biosynthesis (**Figure 4**). It is well known that polyamine biosynthesis is critically regulated in a cell. This class of compounds is known to promote rapid proliferation by specifically contributing to the purine/pyrimidine biosynthesis in a cell and is thus considered a marker for neoplastic progression (44). Thus, the fact that our predictive metabolomics analysis shows this class of metabolites to be significantly deregulated in the pre-cancerous stage (prior to when invasive tumors starts proliferating rapidly) is not surprising. This was further validated upon analysis of serum from KPC mice – mice that did not have observable tumors, but had PanINs, showed an increased serum polyamine concentration (**Figure 4D,E**). Interestingly, upon comparing the abundance of top 5 species as the KPC tumors progressed, we observed that the bacterial species, *Lactobacillus reuteri*, was detected in the 4-month sample, while it was below detection in the earlier time points (**Figure 5**). In our metabolomic reconstruction analysis, *Lactobacillus reuteri* was associated with polyamine metabolism. It is reported that in gastric cancer *H.pylori* promotes DNA damage and neoplastic progression by deregulating polyamine metabolism and promoting oxidative stress in the gastric epithelial lining (45–47). While members of *Lactobacillus sp.* are known to affect polyamine metabolism and growth inhibition in gastric cancer (46), their role in pancreatic cancer remain undefined. Consistent with this, we identified serum polyamine concentration was significantly higher in 4 month KPC samples (without observable tumors but with advanced PanINs) compared to the 2-month samples (**Figure 4D**).

In summary, early detection of PDAC has been an extremely challenging task. In this context, our results show for the first time that microbial dysbiosis and its altered metabolic pathways may potentially be exploited to develop a non-invasive biomarker panel. Further, our results indicate a definitive shift in the microbial composition sufficiently early in the tumor development timeline. Thus, a comprehensive fecal analysis for the change in the bacterial species (as shown in **Supplementary Figure 4**), along with a serum analysis for pan-polyamines, can be further evaluated in the patient-derived samples.

While based on spontaneous mouse models for PDAC, our study forms the first step towards understanding how the microbial dysbiosis during tumorigenesis can play a role in the metabolic regulation of active proliferation of tumor cells – by regulating polyamine metabolism and influencing purine/pyrimidine biosynthesis.

## Methods

### Ethics Statement

All animal studies were performed according to the protocols approved by IACUC at University of Miami, USA (#16-066) in accordance with the principles of the Declaration of Helsinki. Serum from de-identified pancreatic cancer patients and healthy controls were obtained according to the approval from the IRB at University of Minnesota (1403M48826) accorded to Dr. Sulagna Banerjee as an author of the IRB proposal when she was in the Faculty of that University. All authors had access to all data and have reviewed and approved the final manuscript.

### Animal models and experimental design

Spontaneous pancreatic animal model KRAS^G12D^ TP53^R172H^Pdx-Cre (KPC) animals from both genders were enrolled in the study at 1 month of age. PDX-cre (Cre) animals that were age matched with the KPC animals were used as control. Fecal samples were collected at 2 months; 3 months and 4 months of age. Initially, 8-9 animals were kept in each group. At 4 months, all animals were sacrificed according to protocols approved by University of Miami Animal Care Committee. Gut and pancreas samples were flash frozen in liquid nitrogen. Pancreas tissues were formalin fixed for paraffin embedding and histochemical analysis. Blood was collected by cardiac puncture prior to euthanizing the animals. Plasma and serum samples were stored for analysis of polyamines.

### Isolation of DNA

DNA from the mouse fecal samples was isolated using the Power Soil DNA Isolation Kit (Qiagen) according to manufacturer’s instructions. All samples were quantified using the Qubit^®^ Quant-iT dsDNA High-Sensitivity Kit (Invitrogen, Life Technologies, Grand Island, NY) to ensure that they met minimum concentration and mass of DNA and were submitted to University of Minnesota Genomics Center for further analysis by 16s amplicon sequencing or Whole Genome Shotgun sequencing (WGS).

### 16s Pyrosequencing and microbiome analysis

To enrich the sample for the bacterial 16S V5-V6 rDNA region, DNA was amplified utilizing fusion primers designed against the surrounding conserved regions which were tailed with sequences to incorporate Illumina (San Diego, CA) flow cell adapters and indexing barcodes. Each sample was PCR amplified with two differently bar coded V5-V6 fusion primers and advanced for pooling and sequencing. For each sample, amplified products were concentrated using a solid-phase reversible immobilization method for the purification of PCR products and quantified by electrophoresis using an Agilent 2100 Bioanalyzer^®^. The pooled 16S V5-V6 enriched, amplified, barcoded samples were loaded into the MiSeq^®^ reagent cartridge, and then onto the instrument along with the flow cell. After cluster formation on the MiSeq instrument, the amplicons were sequenced for 250 cycles with custom primers designed for paired-end sequencing.

Using QIIME 1.9.2 (Quantitative Insights into Microbial Ecology, version 1.9.2)(48), sequences were quality filtered and de-multiplexed using exact matches to the supplied DNA barcodes and primers. Resulting sequences were then searched against the Greengenes reference database of 16S sequences, clustered at 97% by uclust (closed-reference OTU picking) to obtain phylogenetic identities. Analysis for alpha- and beta-diversity was done with standardized QIIME workflow or the ‘R’ statistical package, as we have shown before (42).

### Metagenomic Sequencing and Microbiome analysis

Shotgun metagenomic library was constructed with the Nextera DNA sample preparation kit (Illumina, San Diego, CA), as per manufacturer’s specification. Barcoding indices were inserted using Nextera indexing kit (illumina). Products were purified using Agencourt AMpure XP kit (Beckman Coulter, Brea, CA) and pooled for sequencing. Samples were sequenced using MiSeq reagent kit V2 (Illumina).

Raw sequences were sorted using assigned barcodes and cleaned up before analysis (barcodes removed and sequences above a quality score, Q≥30 taken forward for analyses). For assembly and annotation of sequences, MetAMOS (49) pipeline or Partek Flow software (Partek^®^ Flow^®^, Partek Inc., St. Louis, MO) were used. These softwares provide powerful tools to filter unique hits between human and mouse-specific genes versus microbial signatures. Alpha and Beta diversity calculations were done using embedded programs within the metagenomic pipeline, or using Stata15 (StataCorp LLC, College Station, TX) or EXPLICET software(50).

Functional profiling was performed using HUMAnN2-0.11.1(33) with Uniref50 database to implement KEGG orthologies.

### Immunohistochemistry

For histochemistry, 4 μm paraffin tissue sections were deparaffinized in xylene and rehydrated through graded ethanol. Hematoxylin and Eosin (H&E) staining were conducted to confirm histological features. Histological scoring was done by an arbitrary scoring method in which scores of 0-4 were assigned based on the dysplasia observed in the tissues. A score of 0 was assigned to the histologically normal pancreas while a score of 4 was assigned to a fully dysplastic tissue (that observed in the late stages of tumor development in the KPC model). All scoring was done in a blinded manner by 3 separate investigators.

### Polyamine estimation

Polyamine was estimated from the serum of KPC animals between the age groups of 2-8 months. The animals of 1-2 months old showed normal pancreas histolology, those 3-4 months old showed various degrees of PanINs and those that were 4-8 months old had observable tumors. Polyamine was estimated by Total Polyamine Estimation Kit (Biovision, Milpitas, CA) according to manufacturers’ instruction after diluting the serum 1:25 in the assay buffer.

### Statistical Analyses

#### Microbiome analysis with QIIME and whole genome analysis pipelines

OTU tables were rarefied to the sample containing the lowest number of sequences in each analysis. QIIME 1.9.2 was used to calculate alpha diversity (alpha_rarefaction.py) and to summarize taxa (summarize_taxa_through_plots.py). Principal Coordinate Analysis was done within this program using observation ID level. The Adonis test was utilized for finding significant whole microbiome differences among discrete categorical or continuous variables with randomization/Monte Carlo permutation test (with Bonferroni correction). The fraction of permutations with greater distinction among categories (larger cross-category differences) than that observed with the non-permuted data was reported as the p-value. Relative abundance of species identified through the metagenomic pipelines were compared using non-parametric Mann-Whitney U test at p<0.05 after FDR correction. Apart from statistical functions embedded within the metagenomic pipeline mentioned above, we have used GraphPad Prism (GraphPad Inc, La Jolla, CA) or Stata15 (StataCorp LLC, College Station, TX) for different statistical analyses, mentioned in the text and figure legend as appropriate.

### Data Availability

Microbiome raw data sequences are available from EMBL ArrayExpress with the accession number E-MTAB-6921.

## Supporting Information

**Supplimentary Figure 1**: Several OTUs mapping to the genus Bacteroides showing increased or decreased relative abundance between 1 and 6 months old KPC microbiome. Overall output shows “no change”. Hence, deeper analysis of microbial species, with higher species-level identification is the only way to decipher the actual role of microbiota in host health and disease.

**Supplementary Figure 2**: Heatmap of bacterial classes at two months age in ‘Cre’ and ‘KPC’ gut microbiome. There is an even spread of the classes within the two genotypes.

**Supplementary Figure 3**: Heatmap of bacterial species in 4 months old ‘Cre’ and ‘KPC’ mice with major changes (both up and down) in many species. The differences have been further elaborated in other figures and the manuscript text.

**Supplementary Figure 4**: Significantly changed species between 2 months old and 4 months old KPC mice microbiome. (A) A heatmap of 82 species found to be significantly changed. Species marked with an arrow had significantly diminished relative abundance at 4 months age, while the others had expanded relative abundance. (B) All 82 species represented 14 different phyla, with dominant representation from Proteobacteria, followed by Firmicutes.

Author contributions
SuB and SaB conceptualized the work, drafted the manuscript and obtained funding for this work, RM, KK, NA and LDM acquired the data, SaB, SuB, RM and KK analyzed the data, SuB, SaB, FM and NM critically reviewed the manuscript.

